# *Bio-*informatics: Integrate negative controls to get the good data

**DOI:** 10.1101/2024.10.08.617225

**Authors:** Rob van Nues

## Abstract

High-throughput datasets, like any experimental output, can be full of noise. Negative controls, i.e. mock experiments not providing information concerning the biological system under study, visualise background. Overlooking this ‘training set’ of wrong examples in publicly available datasets can seriously undermine validity of bioinformatics analyses. We present a program, COALISPR, for explicit and transparent application of negative control data in the comparison of high-throughput sequencing results. This yields mapping coordinates that guide fast counting of reads, bypassing the need for a reference file, and is especially relevant when small RNA sequencing libraries contaminated with breakdown products are analysed for poorly annotated organisms.

We have re-analysed small RNA datasets for mouse and fungus *Cryptococcus neoformans*, leading to consistent identification of miRNAs and of fungal transcripts targeted by siRNAs. Cryptococcal Argonautes are directed to spliced transcripts indicating that RNAi must be triggered by events downstream of intron removal. Negative control datasets contain large amounts of ribosomal RNA (rRNA) fragments (rRFs). These differ from small RNAs associated with RNAi, making a biological role for rRFs in association with Argonautes unlikely. Background signals enabled identification of cryptococcal genes for RNase P, U1 snRNA, 37 H/ACA and 63 Box C/D snoRNAs, including U3 and U14 essential for pre-rRNA processing. To gain meaning, high-throughput RNA-Seq analyses need to incorporate negative data.

**GRAPHICAL ABSTRACT:** 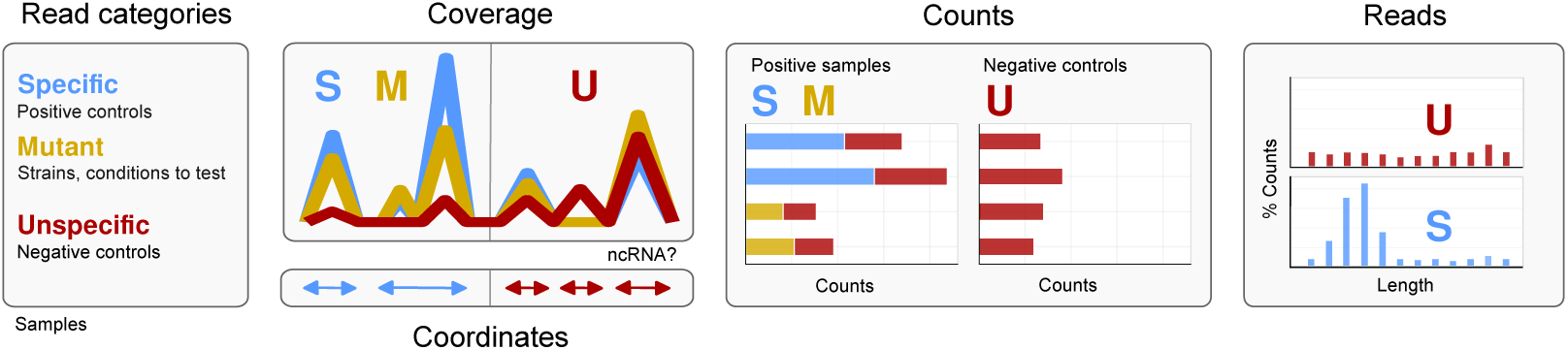

## INTRODUCTION

An ongoing problem in bioinformatics analysis of high-throughput data is that of distinguishing meaningful information against a backdrop of noise and accidental peaks which, by their omnipresence in repeats of experiments with various substrates, appear to be linked to methodology rather than inputs (1). This problem further extends to meta-analyses of various independent datasets if this complication is not recognized. Sound statistical underpinning does not validate conclusions about signals when experimental methodologies that address the removal of noise have not been adhered to in the data analysis. This paper describes how background can be used in the interpretation of high-throughput sequencing results.

Current day bioinformatics tools for analysing high-throughput sequencing datasets are aimed at outputs obtained for protein-gene transcripts. Transcript counting, for example, relies on description of genes, annotated in GTF or GFF reference files regarding coding sequences (CDS), untranslated regions (UTRs), introns and exons. Descriptions for non-coding RNA (ncRNA) genes are often absent in annotation files and their transcripts are not counted. In this context, ncRNAs are treated as ‘background’ although experimentally they tend to interfere with signal-to-noise ratios and effective preparation of sequencing libraries. For this reason, highly-conserved and therefore well-detectable ncRNA molecules, such as ribosomal RNA (rRNA) or transfer RNA (tRNA), are often removed from the input by means of depletion kits or from the data by sequential mapping of reads. This filtering needs to be deduced from the Materials and Methods of experimental papers. How these ‘tools of the trade’ carry over to proper usage of RNA-sequencing (RNA-Seq) datasets deposited in public repositories like the Gene Expression Omnibus (GEO) can, however, remain unclear, especially when in the experimental field the relevance of particular control datasets is still in need of explanation (1).

The ‘controls’ are test-reactions within an experiment molecular biologists rely on to reach valid conclusions. Negative (or mock) controls are experiments that give an outcome without biological significance. They are essential for determining the noise within an experiment. The common practice of using negative controls extends to analysis of high-throughput RNA-Seq data which is not always recognized and has led to major misinterpretation of available data for small ncRNAs. In small RNA-Seq, mock controls help to identify breakdown products of longer molecules that contaminate the datasets (2). Here, we propose standard inclusion of negative control data in initial bioinformatics analyses to evaluate libraries in a dataset. This also helps to highlight characteristics of datasets that affect their comparability.

It may be thought that all RNA as a product of gene-transcription belongs to one category, thereby posing the same problems to solve when it comes down to bioinformatics. This turns out to be too simplistic a view given the plethora of high-throughput data that can be generated. To reduce experimental noise, RNA input for sequencing is often obtained by selective methods such as size-selection, RNA immunoprecipitation (RIP) by means of epitopes on RNA binding proteins (RBP), or variants thereof involving UV crosslinking (3, 4). These methods enrich for a particular subset of all the RNA available in a cell. Then, the prevalent practice of library comparison using applications designed to measure differential gene expression, e.g. Deseq2 (5) or EdgeR (6) loses applicability because the internal standards formed by an unaltered population of transcripts and required for normalization by these tools go missing. The link with transcription is also severed when the biological process generating the RNA follows a different route. During RNA interference (RNAi) in various model organisms, amplification of siRNAs results from reverse transcription of the target RNA by RNA-dependent RNA-polymerase (RDP), followed by dicing the double-stranded product and final association of single-stranded small RNAs with Argonaute proteins (7, 8). Stress encountered by cells can trigger an onset of RNAi: After strain construction by site-directed mutagenesis with CRISPR-Cas9 (9), siRNAs were found against exogenous components transformed into the cells and this varied among biological replicates (RvN, Elizabeth H. Bayne unpublished).

In view of the variability in the kind of datasets analysed, it is relevant to know what count as ‘specific’ and what as ‘background’ (noise). For a better understanding of the information datasets convey, it would therefore be advisable to make this distinction explicit. This would also help to be able to check the extent in which extraneous nucleic acids (e.g incorporated during strain construction) pervade the RNA-seq data. In order to do this, we have developed a Python application, COALISPR (*CO*unt *ALI*gned *SP*ecified *R*eads), to systematically clean up datasets by transparent application of control data. By comparing these to experimental data, coordinates for mapping regions are obtained that guide fast counting of reads, bypassing the need for a reference file at this stage.

To demonstrate the validity of our approach we re-analysed RNA-seq datasets for mouse (2), baker’s yeast (10–12), and the fungus *Cryptococcus neoformans* (13, 14), an opportunistic pathogen. We noticed that siRNAs in this fungus were directed to transcripts that have undergone splicing so that RNAi must be triggered by events downstream, contrary to what has been published (13). With the output of COALISPR We show, using ribosomal RNA fragments (rRFs) as an example, that novel insights cannot solely be based on large numbers that appear to promise statistical significance: As a major part of negative control data, rRFs do not bind specifically to Argonaute proteins. By exploring signals specific for negative control data, over 120 ncRNAs not previously annotated for *Cryptococcus* were identified, which was corroborated by phylogenetic analyses.

## MATERIALS AND METHODS

### Software

All figures (but Figures 7B-D) were adapted from originals generated with COALISPR as described in detail in the documentation available at https://coalispr.codeberg.page/. A how-to guide and tutorials explain each step from raw sequence to output. COALISPR is accessible from https://pypi.org/project/coalispr/ and https://codeberg.org/coalispr/. The program is written in PYTHON (https://www.python.org) and depends on software libraries PANDAS (15) (https://pypi.org/project/pandas/), NUMPY (16)(https://pypi.org/project/numpy/), PYSAM https://github.com/pysam-developers/pysam), MATPLOTLIB (17) (https://pypi.org/project/matplotlib/), and SEABORN (18) (https://pypi.org/project/seaborn). Input files were obtained with the STAR (19) aligner (https://github.com/alexdobin/STAR) and, for collapsing reads, PYCRAC (20) (https://pypi.org/project/pyCRAC/). All figures were made with INKSCAPE (https://inkscape.org). Genome browsers IGB (21) (https://bioviz.org) and IGV (22) (https://igv.org) were used to scan reads at 1 nt resolution. Software development and all testing was done on a Clevo N750HU (from PC Specialist, https://www.pcspecialist.co.uk/) with an Intel Core i7-7700HQ CPU @ 2.80 GHz and 32 GB RAM, running Slackware linux 64bit (http://www.slackware.com/) with above software obtained via https://slackbuilds.org.

### Datasets

Various datasets have been used to demonstrate application and use of negative controls. The mouse biochemical analysis and miRNA sequencing were described in Sarshad et. al. (2). The miRNA-Seq data have been kindly provided by Markus Hafner. Baker’s yeast data had been used in Sharma et al. (11), van Nues et al. (10), and Gerhardy et al. (12). The siRNA- and RNA-Seq data for *C. neoformans* were from Dumesic et al. (13) Burke et al. (14) and Wallace et al. (23). Where possible, raw sequences were retrieved from the given GEO depositories and re-analyzed by aligning with STAR (19) directly or after collapsing with PYFASTQDUPLICATEREMOVER.PY from the PYCRAC (20) package. Generated bedgraph and bam files were used as input for display and counting, respectively, and have been deposited at https://zenodo.org (<u>DOI</u> https://10.5281/zenodo.12822544) for mouse and *C. neoformans* together with files that would enable a starting point for running COALISPR scripts.

### Annotation files

Annotation files used for the re-analysis of published datasets were prepared as described in the tutorials of the COALISPR documentation. Annotations for mouse were from release M30 for the GRCm39 genome as described (2); miRNA sequence annotations and their targets were extracted from the miRNA database, miRDB_v6.0_prediction_result.txt.gz (24) with the help of gene2ensembl.gz downloaded from https://ftp.ncbi.nlm.nih.gov/gene/DATA/. Yeast reference data has been as described (10). Analysis of *C. neoformans* data was based on release 55 of GCA_000149245.CNA3 from https://ftp.ebi.ac.uk/ensemblgenomes. Annotations were derived from H99.10p.aATGcorrected.longestmRNA.2019-05-15.RiboCode.WithStart.gtf (23) and, via https://ftp://ftp.ncbi.nlm.nih.gov/, genbank file GCA_000149245.3_CNA3_genomic.gbff. For Supplementary Figure 3, annotations from Wallace et al. (23) were compared to the original GFF descriptions used by Janbon et al. (25) downloaded from https://genome.jgi.doe.gov/. All ncRNAs that were not yet or incorrectly annotated were found by scanning traces in genome browsers IGB (21) and IGV (22).

### Phylogenetic Analysis

Sequences with significant hits in all siRNA and mRNA samples that were not annotated (13, 14, 23, 26) were checked for a correspondence at RNAcentral (https://rnacentral.org/sequence-search/) and Rfam (https://rfam.org/family/). Orthologous ncRNAs in *Tremellomycetes* were found by means of iterative searching by NCBI BLAST (https://blast.ncbi.nlm.nih.gov/Blast.cgi) as described previously (11, 27). Known snoRNAs with comparable guides were identified via the SNORNA ORTHOLOGICAL GENE DATABASE (28) (http://snoopy.med.miyazaki-u.ac.jp/). Alignments were prepared with JALVIEW (https://jalview.org), and exported in the Stockholm format for building consensus secondary structures with R2R (29) (https://sourceforge.net/projects/weinberg-r2r/) and for publication alongside images in the documentation for Coalispr.

### Analysis of libraries with COALISPR

#### Principles

COALISPR consists of a collection of command-line scripts for quick, selective comparison and visualisation of high throughput sequencing data. The program dissects the data per chromosome using bedgraphs for input and helps to retrieve read counts from associated bam files. For relatively small fungal genomes, it can display over 100 bedgraphs in one panel.

The program mimics visual subtraction between positive and negative control samples referred to below, but acts on bedgraph traces, as illustrated in the schematic of Figure 1. Each bedgraph trace stands for one strand to which reads in a sample (a library of a dataset) are mapped to. For each sample, stranded bedgraph data from files output by aligners like STAR (19) are tabulated per chromosome by splitting and summing over bins. Bins have a defined size that can be set in configuration files to, say, 50 bp. Chromosomes differ in length and are kept apart to facilitate building displays with MATPLOTLIB (17). The binning and further processing are done with PANDAS (15).

**Figure 1.**
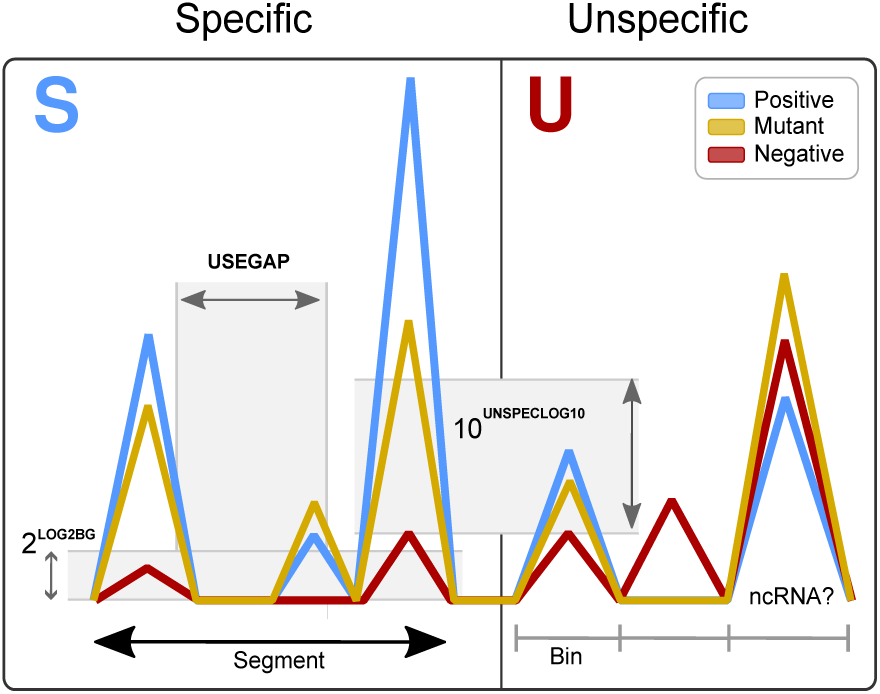
Identification of segments with s*pecific* reads (based on positive controls) depends on the setting of indicated parameters (bold, capitalised). **UNSPECLOG10** sets a threshold for reads with some overlap to reads in the negative control. The demarcation of read segments (clusters with contiguous reads) relies on **LOG2BG** and **USEGAP** that bridges peaks remaining after filtering *unspecific* reads. Obvious peaks shared by all samples could indicate a ncRNA.

Tabulated bin values are then compared between negative and positive samples. When a bin contains hits in a negative control, while comparable values are found for other samples, the reads in that bin are taken to represent a background signal. These reads are dubbed ‘unspecific’ (U) and can build a low signal, but when they derive from very abundant molecules like tRNA or rRNA, they accumulate and form substantial peaks in the bedgraph traces. Therefore, in order to distinguish such high levels of noise linked to the applied methods from signals specifically detected by the experiment, comparison with negative control data is key.

Specific signals (S) are marked by bins that mainly contain reads in the case of the positive controls or mutant samples. These bins show coverage for reads representing RNAs that can be linked to a condition under study or the protein they have been co-purified with. These reads are (mostly) absent from negative controls, so that the difference in coverage rather than signal intensities is used as the leading parameter to specify reads as specific or unspecific. While the difference in coverage is the major determinant, various configuration settings help to define segments with specific signals (the remainder being unspecific). With parameter **LOG2BG** a base level is set below which signals are not taken into account. The second parameter, **UNSPECLOG10**, sets a threshold when reads of positive controls and those in negative controls overlap; it indicates the minimal acceptable fold-difference between specific and unspecific signals. Then, to account for missed or dismissed data, the tolerated gap between genomic peak coordinates can be set with the constant **USEGAP** to define a contiguous segment for counting reads. These settings are part of configuration files that need to be edited from templates to fit the specifics of each dataset that is to be evaluated.

Comparison of read signals collected in bins, instead of directly per nucleotide, is a pragmatic decision. It enables tabulation of all data once imported and thereby the application of vectorisation techniques to process all samples in one go. Re-organising bedgraph signals to a common index, i.e. bin-related coordinates, reduces the resolution but is the pivot of the approach. By concentrating reads, binning improves definition of difference trends because read-coverage variations between samples will have less impact. The reduction in resolution helps, apart from decreasing memory usage, to distinguish genome regions with specific signals from those with unspecific, background peaks. This read sorting, preceding subsequent counting and other analysis steps, is the main, novel contribution of COALISPR to current bioinformatics and highly relevant when size-selection steps cannot reduce contaminating signals, as is the case for small RNA-Seq. An example of sorting reads into the two categories is shown in Figure 2. Note that COALISPR is not intended to replace genome browsers (21, 22) but for complementary, parallel use.

**Figure 2.**
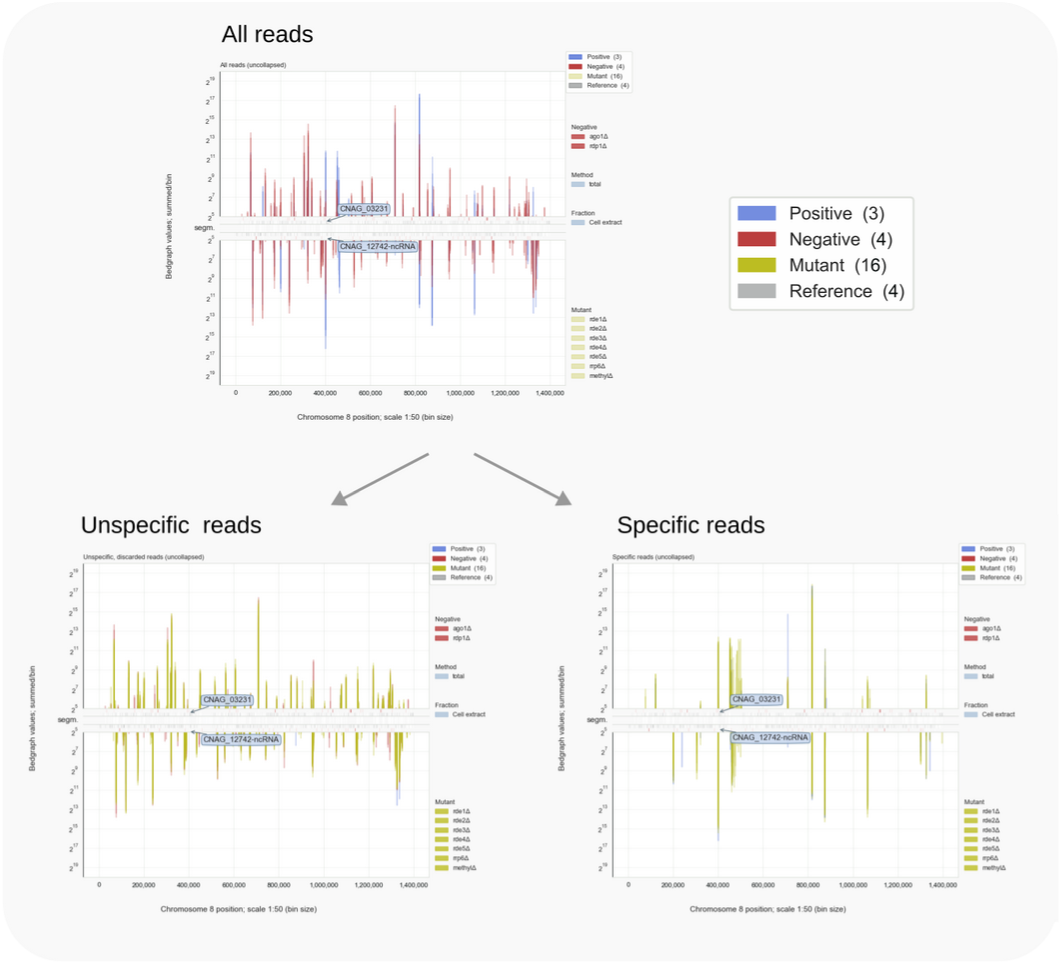
Specifying reads for chromosome 8 of *C. neoformans* H99 by comparing signals for mutants, positive and negative controls. Parameters **UNSPECLOG10** = 0.78, demanding at least a ∼6-fold difference between specific signals and that of background in the same bin of 50 bp; **LOG2BG** = 5. For the top panel, traces for the mutants were set to invisible.

#### Counting reads without an annotations file

Apart from visual comparison of data-traces, the program has been developed to obtain count-data for small non-coding RNAs. Genome annotation files for most organisms lack information on such RNAs, while programs like HTSEQ (30) or PYCRAC (20) base counts on annotated features. Thus, to count small RNAs with these programs an annotation file has to be prepared from scratch. This is laborious, error-prone, and not very scalable to other organisms or when a major update of the genome is published. By incorporating negative control data in the analysis, this step can be evaded/postponed. The bin comparison applied by COALISPR not only helps to distinguish meaningful reads but also to systematically obtain read counts for various kinds of data.

The separation of reads into those that fit the positive controls and those that are common to all samples (i.e. overlap with negative controls) defines the chromosomal regions belonging to the either of the two read categories (Figure 1). Counting of reads can thus be based solely on genomic coordinates in bam alignment files the bedgraphs derive from, omitting any check for an overlap with an annotation reference. The only requirement is that bam files have been sorted by coordinate. The program implements this principle and supports fast counting of reads when these have been collapsed prior to the mapping. Collapsing with PYFASTQDUPLICATEREMOVER.PY (20) takes all identical sequences as one, but tracks the total number that represent a cDNA, which enables counting of all identical reads at once. Various count data can be visualised by means of scripts that are built around SEABORN (18) and MATPLOTLIB (17), for example for overall read numbers (Figure 3).

**Figure 3.**
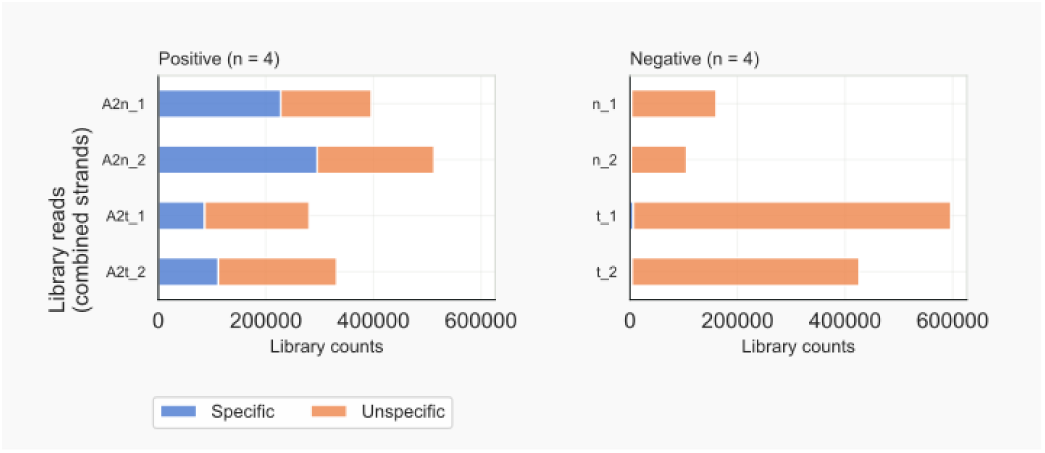
Counts for specified reads in mouse miRNA libraries. Large fractions of positive control reads are unspecific and shared with the negative controls; **UNSPECLOG10** = 0.905; **LOG2BG** = 4.

Apart from counting overall numbers (Figure 3), the program can give information on counts and characteristics for reads that have been mapped to a specific locus. A telling characteristic for reads is their length, which would relate to the RNA binding pocket of a RBP, while the starting nucleotide can be a telltale for small RNAs associated with Argonaute proteins involved in RNAi. For this reason, length-distributions for reads can be generated for each library, a particular region or for the whole dataset, whereby specific and unspecific reads are distinguished from each other.

#### Annotate count outputs

Reference files can be used in the program for annotating known genetic elements when displaying bedgraph traces (Figure 2). The same annotation files can be scanned for identifying regions with reads in count files, which can be sorted according to value, providing an overview of the most abundant molecules in a dataset.

## RESULTS AND DISCUSSION

The crux of our approach is combining essential biological methodology, the BIO, with bioinformatics by integrating experimental control data into the mathematical analysis. Experimenters do not just generate data, they produce data and control data; the latter are key to draw valid conclusions from the former. What is meant by control data?

### Control data

Biological experiments encompass approaches that historically precede but are conceptually similar to machine-learning methods when training is done on ‘bad’ vs ‘good’ examples in the data.

Experimental success lies in the pragmatic possibility to make a proper assessment. That is, from the output of the experiment a conclusion concerning some hypothesis on a biological system can be drawn. Experimental results can only be meaningful if standards are present within the experiment to evaluate the overall output against. This because a biological experiment involves components that are not fail-safe. Reagents can expire and no longer be active. Other problems relate to methodology. For example, in small RNA-Seq the short molecules isolated are always mixed with remnants of longer molecules. A size-selection step during the RNA and/or cDNA library preparation will not resolve this difficulty. In general, RNA-Seq libraries will contain contaminants because of electrostatic interactions between nucleophilic molecules in the starting input or when these molecules have some affinity to materials used during preparative steps. To address such issues biological experiments contain internal tests for proving that outcomes obtained with the followed procedures would have been valid.

The standards that render the internal tests are called the positive and negative controls. The negative control is a sample that is not expected to provide an informative answer; also called ‘mock’ control, it shows the noise in the experiment. In contrast, a positive control should give output that is specific, i.e. adheres to the current knowledge available for the biological system under study. It tells that the experimental conditions produced a useful answer. To assess the effect of a mutation (or another condition) is to check what happens to the specific output relative to that of the controls.

The noise of the negative control, the meaningless signals that are also expected for other samples in the same experiment, can be seen as the ‘bad’ examples in a machine-learning training set. Such noise is visible as a diffuse smear in a biochemical analysis of RNA used for the construction of cDNA libraries in RNA-Seq. For example, in a study of microRNAs by Sarshad et al. (2) the gel-image of <u>their</u> Figure 3D (or 3F) shows radioactively labelled RNA that is representative for RNA-Seq input. The RNA was purified from cells that only produce one functional Argonaute protein (AGO2) after induction of the gene expressing an epitope-tagged variant (+Dox). The protein was isolated by immunoprecipitation with the Flag antibody (IP:Flag) from two cellular fractions (cytoplasmic, C and nuclear, N). The same procedure was done with cells that were not induced (-Dox) and therefore not expected to make AGO2. In this experiment, the -Dox samples are the negative control to ensure that output obtained for the samples of real interest (+Dox) can be trusted. The latter samples could be dubbed the positive controls because the experimental outcome showed the co-purification of AGO2 with a size-restricted population of RNA molecules (the fat black band at the level of ‘:F’ in the label ‘IP:Flag’). This was in agreement with the methodology and published findings that miRNAs of ∼22 nt bind to Argonaute proteins in mammals. Other controls in this example are for correct fractionation into cytoplasmic components (CALR) and nuclei (LMNA).

The absence of tagged protein in the -Dox samples demonstrated that the gene was not expressed and indicated properly working induction and immunoprecipitation procedures. Visual subtraction leads to the conclusion that only the fat band in the lanes of the positive samples represents the meaningful outcome of the experiment: the signals common to all lanes, the background smear of randomly broken molecules present in the extract after smashing up the cells, can be ignored. But not for the follow-up experiment. The RNA smear, visible in the -Dox lanes, is still present in the positive samples and will be part of the input for RNA-Seq, which was subsequently done on all four samples (2) to enable a similar comparison with respect to the sequencing results. Such comparison with appropriate negative controls, that remains often implicit or seems overlooked, is automated by COALISPR.

### High-throughput datasets can be full of noise

In the discussion of their results, Sarshad et al. (2) only mention the relevant data, the miRNAs isolated from the +Dox samples. Still, the -Dox dataset would have been used to determine which sequencing reads could be ignored in the +Dox data, enabling identification of sequences that specifically associate to AGO2. This filtering is common practice and hardly mentioned or properly explained in publications, which impacts reproducibility. In other words, this essential step can easily be overlooked, especially by non-experimenters, and is possibly the reason negative datasets are not taken into account by at least some meta-studies. This is one of the problems we discuss in this paper, and which we have addressed by developing a Python application to systematically clean up and count (small) RNA-Seq results.

The strategy of ruling out ‘bad’ examples as in a machine-learning training set is evident when experimenters remark that some particular hits were not used or filtered from the dataset by sequential mapping to sequences of known ncRNAs. Statements concerning omitted reads are indicative for the fact that high-throughput data always contain sequences different from those that are deemed relevant for the system under study.

When purifying reagents, which determines the success of an experiment, irrelevant molecules can be fished out from the input-sample. This will minimise contaminating signals in the eventual output. Depletion of rRNA (31) makes the RNA-Seq more efficient by improving the coverage for data the researchers are interested in. Other experimental approaches are abound with a similar aim of improving the signal to noise ratio. For example, as described above, selective purification of an RBP and then sequencing the associated RNA helps to narrow down the population of RNA molecules the protein interacts with. Cross-linking techniques have been developed to enrich even more for biologically-relevant over spurious molecular interactions, be it between proteins and RNA (3, 4) or between RNA-molecules bound to a protein of interest (32).

The words ‘narrow down’ and ‘enrich’ are necessary: noise, although reduced, will remain as exemplified by (vague) background bands in gel-images that represent experimental results. Both cDNA library construction and the sequencing reaction involve PCR amplification steps. This means that the noise will be increased along with the signals. Therefore, as illustrated above in Figure 3, the noise can form a more substantial portion of the RNA-Seq output than expected with input of good quality. Such background corresponds to the category of ‘unlikely’ reads that have significant overlap with peaks in samples extracted from a ‘blank’ control in Figure 2 of Esteban-Serna et al. (1).

The take-home message is that RNA-Seq data are not ‘clean’; they are messy and not usable as is. To draw reliable conclusions from the data, the irrelevant noise has to be put aside. We propose to make this taken-for-granted aspect of the data analysis explicit and to systematically remove the noise by following common biologists’ practice of using negative controls. With COALISPR we demonstrate how this essence of experimentation can be translated to a bioinformatics approach.

### Ribosomal RNA hits

Very abundant cellular complexes like translating ribosomes are a source for fragments of rRNA, tRNA, or highly expressed mRNAs that create unwanted background signals in RNA-Seq data. Most ribosomal fragments (rRFs) originate from very long (precursor) molecules. This section describes the analysis of rRFs in datasets by means of COALISPR.

#### Mouse miRNA data

By dividing reads into specific signals and unspecific noise, a large fraction of reads in AGO2 IPs was found in negative controls (Figure 3). This was to be expected because of the background smear being part of the input for the RNA-Seq libraries (<u>Sarshad et al. 2018,</u> Figure 3D). The characteristics of the grouped reads, however, are very different (Figure 4). The majority of AGO2-associated RNAs are 22-23 nt, conforming to expected lengths of miRNAs, and have an A as the 5’ nt. Unspecific RNAs are more diverse in length, with RNAs of 31-33 nt starting with a G most prominent. What do the rRFs look like in this dataset? Mammalian ribosomal DNA repeats encode 45S pre-ribosomal precursors. Of these rDNA repeats, an 18S unit on chromosome 17 is annotated for the genome used by Sarshad et al. (2). The display of Figure 5 shows traces for nuclear and cytoplasmic negative controls (red) that are indistinguishable from those for the AGO2 IPs (blue) in the case of sequences mapping to rDNA (top). Compare this to loci for miRNA genes, expected to be covered by reads associated with AGO2 rather than those isolated from uninduced samples. This can be seen in the range chr17:18050000-18052000 with a miRNA cluster of Mir99b, Mirlet7e, and Mir125a (ENSMUSG0000207560 was not detected). When the complete 45S rDNA unit was included for mapping, no major change was observed for AGO2-specific reads, while cDNAs derived from rRFs covering the 45S rDNA region were comparable for all samples (data not shown). No group of rRFs or rRFs-derived cDNAs, including those that adhered to the characteristics of miRNAs, could be discerned that stood out in positive samples compared to the negative controls with respect to length and start nucleotide (Figure 6). Thus, while being among the most abundant reads in this dataset (Supplementary Tables, sheet S1), rRFs cannot be proven to be specifically associated with AGO2. This outcome undermines published meta-analyses claiming such a role for rRFs, a suggestion originating from over-optimistic assumptions that input RNA had been sufficiently pure and that background in the form of a smear on gel (<u>Sarshad et al. 2018,</u> Figure 3D) would not produce relevant count numbers in RNA-Seq. This would all have been preventable had this bioinformatics analysis incorporated available negative control data. The conclusion is, that rRFs are not co-purified with mouse AGO2 because of a biologically meaningful interaction.

**Figure 4.**
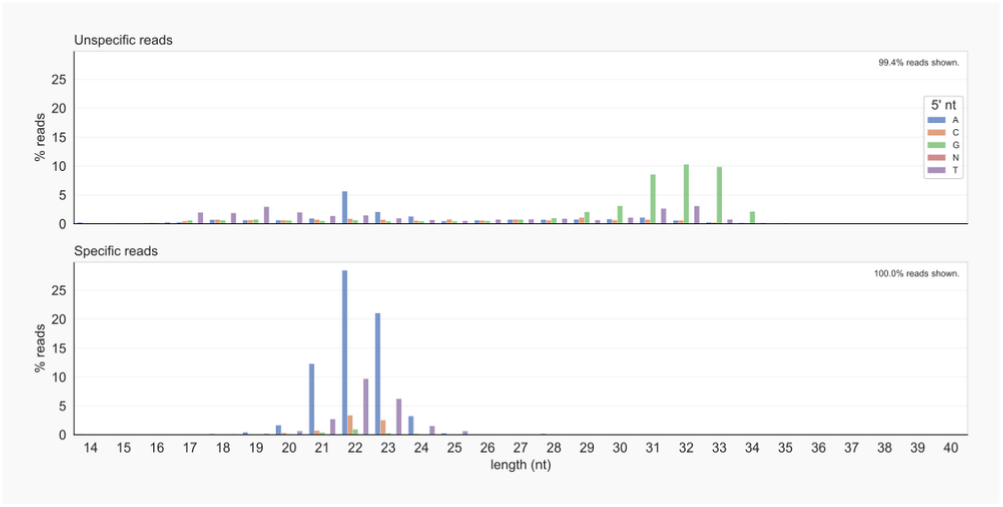
Lengths and start (5’ nt) of specified reads in mouse; RNAs shared with negative controls (top) differ from AGO2-associated RNAs (bottom).

**Figure 5.**
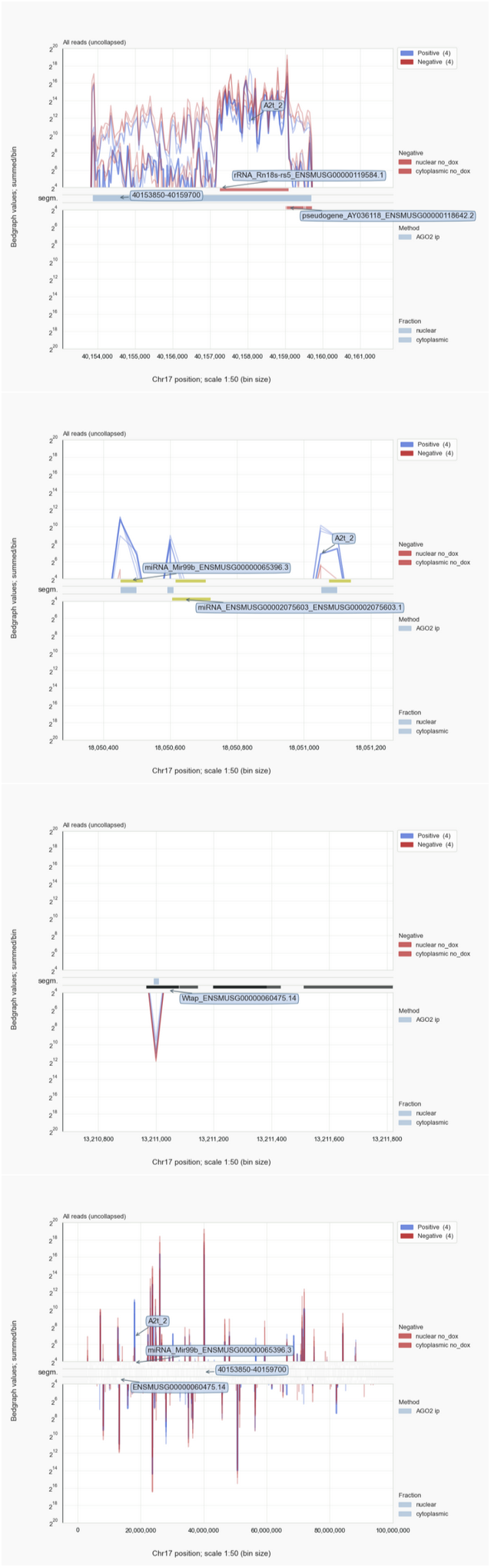
Trace analysis for chromosome 17 in mouse; see text for details; from top (1) to bottom (4): 1: Common 18S rRFs, 2: AGO2-specific miRNA reads, 3: An unspecific hit for Wtap, a predicted target for 154 miRNAs in miRDB, 4: All reads for chromosome 17.

**Figure 6.**
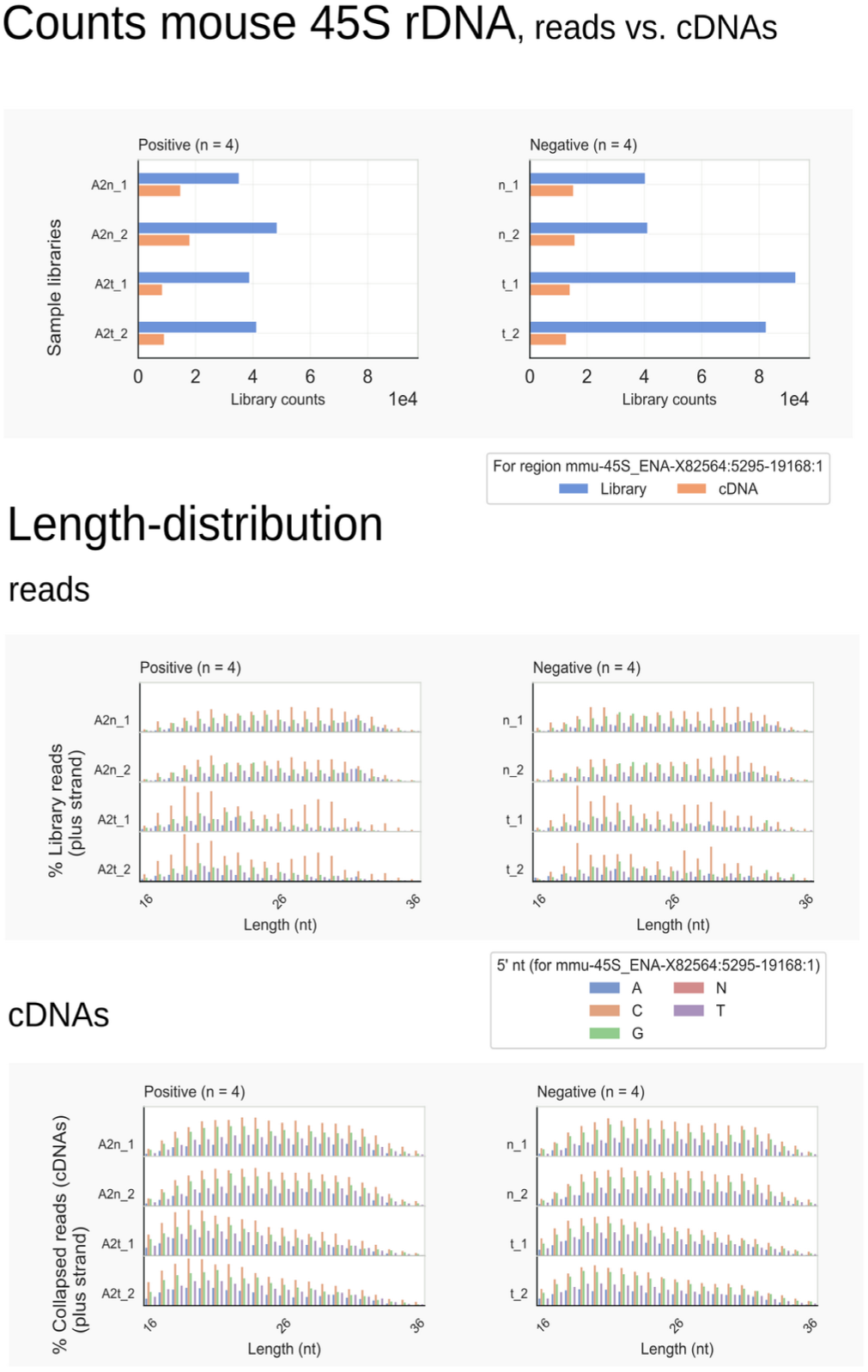
Mouse 45S derived rRF-cDNAs and rRFs. In comparison to negative controls, enrichment of a class of rRFs bound to AGO2 is not supported by counts (top) or length-distributions (bottom).

#### Yeast pre-rRNA processing factors

The finding that rRFs are common to all samples does not mean they cannot represent specific interactions between rRNA and proteins. Techniques involving UV crosslinking of RNA to proteins of interest before purification have been developed for exactly this purpose, identifying binding regions for processing factors involved in ribosome assembly and rRNA maturation (4, 11, 12, 33, 34). Despite the stringent and harsh purification procedures applied in these approaches – whereby the proteins are fully unfolded – background signals from abundant molecules cannot be evaded. Samples that have not been irradiated can be taken as negative controls but in the case of tight complexes or strong interactions this is not ideal. An alternative way to assess noise is by comparing crosslinking data for different, unrelated RNA-binding proteins after multiplexing, that is by processing in parallel with differentiating 5’ bar-coded adapters so that the cDNA libraries could be prepared in the same PCR reaction.

To illustrate how noise can be filtered by using data-sets as mutual controls, various CRAC-samples obtained for processing factors Kre33 (11), Puf6 (12), or termination factor Nab3 (10) were compared (Supplementary Figure 1). Kre33 directs acetylation of cytosine residues in helices 34, and 45 in the 18S rRNA portion of the 90S pre-ribosomal particle and associates with helices 8-10 (11). Especially the latter region stands out from the non-Kre33 traces. Similarly for Puf6, which crosslinks to helices 67-68 of 25S rRNA in the large subunit precursor (12). The more detailed pyCRAC (20) analyses back the specific association of rRFs with Kre33 or Puf6 by the presence of point-deletions of uracil residues in reads that result from UV crosslinking of the RNA to either protein. With this comparative approach relative enrichment of tRNAs and snoRNAs interacting with these proteins could be discerned (11, 12).

These results underline the experiential fact that abundant, spurious rRFs are identified in RNA-Seq libraries despite the stringent purification procedures applied. Any assessment whether rRFs in RNA-Seq data can be linked to a biological role requires proper controls and independent experimental evidence supporting such a function.

#### Fungal siRNAs target rRNA

Most high-throughput analyses catered for by bioinformaticians deal with data obtained for mRNA or RNA fragments that are co-purified with proteins they bind to and rely on extensive gene-annotation files (20, 30). With Coalispr an alternative way for obtaining count-data has been implemented, which is based on genomic coordinates identified by aligning reads to a reference genome. This is the only feasible approach to interpret RNA-Seq data for classes of short RNAs, like siRNAs that have not (or poorly) been annotated. The development and testing of Coalispr has relied on comparison to count-data obtained with HTSeq (30) using manually accumulated reference files for siRNAs found in *Cryptococcus* (13, 14). Compilation of annotations for genomic regions complimentary to these siRNAs provided the insight that siRNAs, contrary to what has been published (13), are raised mostly against transcripts that have undergone splicing (Supplementary Figure 2). Naturally, the possibility that splicing was not entirely complete remains, so that retained introns could be one of many causes for translation failure or leading to messages with premature stop-codons. Such faults, which would be encountered during interaction of spliced transcripts with ribosomes, could turn these transcripts into targets of the RNAi-machinery.

In this respect noteworthy, is the finding of siRNAs associated with Argonaute proteins that are directed against mature rRNA (Supplementary Figure 3). This is seen for *C. neoformans* as well as *C. deneoformans*, but, in the case of the latter, especially after the RNAi response has been restored in cells that did not express their main Argonaute protein, Ago1 (26), or Rdp1 (RvN, Elizabeth H. Bayne data not shown). Another ncRNA associated with ribosomes, SRP RNA, can be targeted by siRNAs (Figure 7, top).

**Figure 7.**
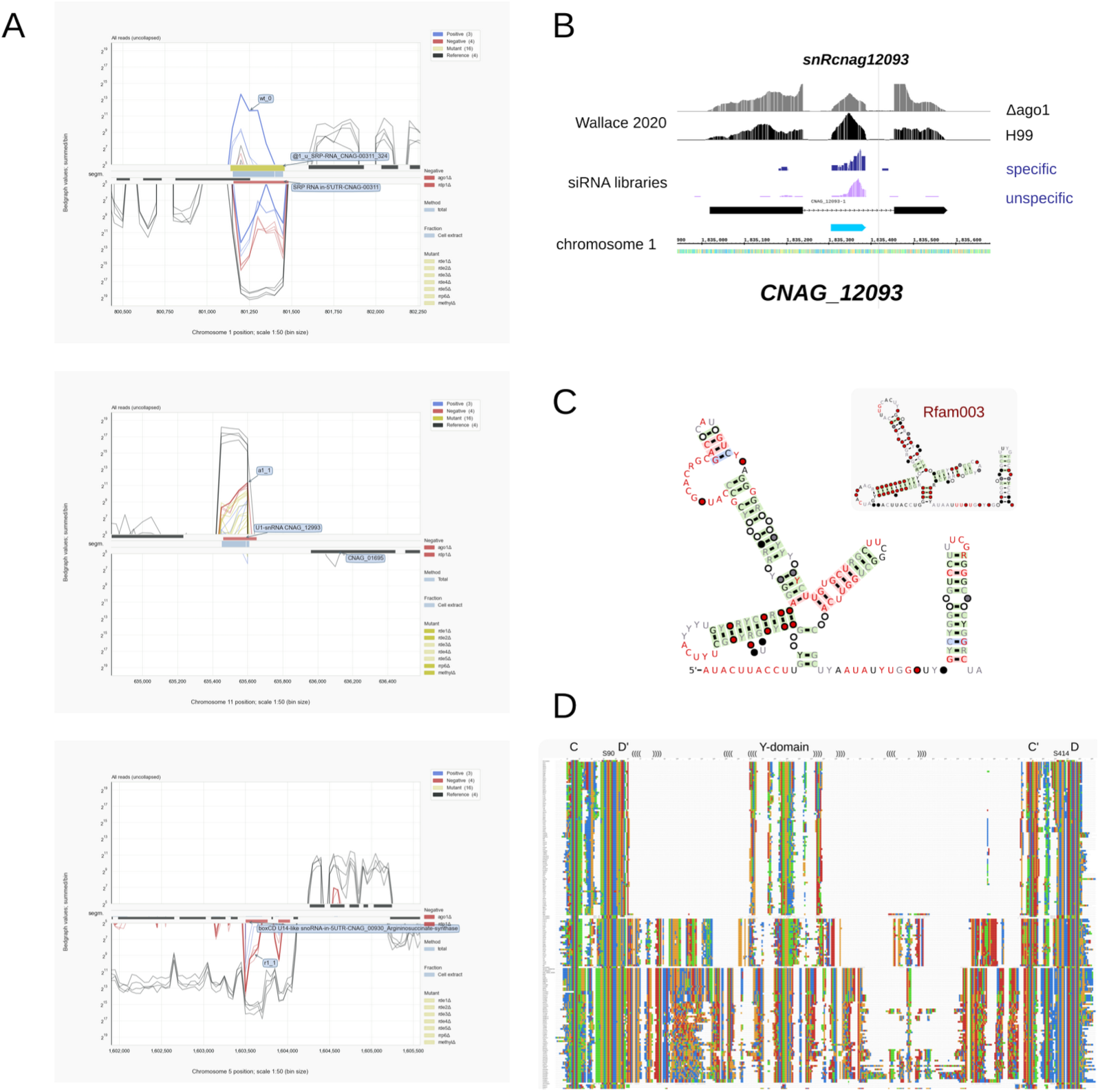
ncRNA genes in *C. neoformans* H99. A: hits for SRP RNA (top), U1 snRNA (CNAG_12993, middle), and snR190 and U14 that derive from introns of *Argininosuccinate synthase* (CNAG_00930) pre-mRNA; mRNA-Seq traces (black) function as reference.; B: IGB display of CNAG_12093 locus with ncRNA expressing an intronic snoRNA. Tracks display reads from RNA-Seq for wild type (H99, specific) and for cells not expressing Ago1 (Δago1, unspecific); C: R2R (29) diagrams of U1 snRNA for *Tremellomycetes* and Rfam003 (inset). The conserved (red) 5’ end interacts with the 5’ splice site GURRGU in *Cryptococcus*; the conserved single stranded region upstream the final stem is the SM-site; D: Alignment of fungal U14 snoRNAs. At the top snR128 of *S. cerevisiae;* snoRNAs from *Tremellomycetes* in the bottom half. Conserved protein binding motifs, C, D’, C’ and D, as well as the Y-domain are indicated above the figure; brackets indicate potential base-pairing.

Comparable to the mouse miRNA and yeast CRAC datasets, large amounts of rRFs were detected in cryptococcal siRNA libraries (Supplementary Figure 3). No reads, however, are specially enriched among rRFs in *Cryptococcus* that would fit the siRNA binding pocket on the fungal Argonaute proteins. We conclude that the observed co-purification of fragments derived from ribosomal RNA transcripts with Argonaute cannot provide evidence for a biological interaction between these molecules.

### Small RNA count data are problematic

The main aim of Coalispr is to sort signals from noise. Peaks in bedgraph traces are specified as specific and unspecific and this process can be followed by comparing the output visually (Figures 2, 5). The specified reads are then counted and these counts, stored in TSV files, can be used for further analyses. Read-counts can be indicative for the quality of libraries and general experimental outcomes, which can be visualised (Figures 3, 4, 6). The main interest, however, will be in comparing the counts between samples for the same RNA in order to understand the biological effect of a specific condition or mutation. For this, read counts need to be normalised, which is problematic for small RNA sequencing data. Why?

Because of the variability in library depth, often related to practical difficulties during preparation and recovery of the cDNA sequencing input, an internal normalisation standard is required. In the case of mRNA sequencing, transcripts from genes are followed and thereby the output of active gene expression. Transcription of genes is highly regulated by transcription factors and depends on the modification status of the C-terminal domains of histones and RNA polymerases and, in many organisms, by chemical modification of the DNA. Gene expression as measured by the levels of mRNA can respond quickly to stress caused by a sudden change in environmental conditions because of RNA-degradation and RNA-RNA interactions (27, 35, 36). In these cases, only gene transcripts involved in biological functions affected by the environmental change under study are supposed to get altered while the majority of gene-expression will not be interfered with. This observation is used by many bioinformatics tools, like DESEQ2 (5) or EDGER (6), to detect relative changes in protein-gene expression. The large set of unaffected, apparent constant transcripts forms an internal standard allowing comparison of RNA-Seq data for cells before and after exposure to an environmental or genetic change.

Quantification approaches of differential gene-expression (DE) by programs like DESEQ2 or EDGER are, however, not applicable to assess changes in read-numbers between libraries that cannot represent the transcriptome. Such libraries are those that have been created after enrichment of RNAs by size or by their association to a particular protein. Small RNA libraries of siRNAs also have this characteristic. Formation of siRNAs, other than miRNAs that are controlled by gene-expression, appears to occur as a response to some change in cellular exposure to harmful nucleic-acids, like invasive viral RNA or transcripts from transposons. Therefore, it can be assumed that a change in siRNA levels could be the result of a stochastic or immediate reaction triggered by conditions peculiar to a specific strain or a group of cells. Observed differences in siRNA populations between biological replicates might be an indication for this (RvN, Elizabeth H. Bayne data not shown). By lack of evidence that siRNA levels are controlled by a well-regulated process like gene expression, there is no internal reference, a base line, that can be decided on. Even if that would be the case, DE-analysis of siRNAs is hampered by, compared to the number of mRNA genes, a relative low frequency of unique loci siRNAs can be mapped to and thus a low number of molecules that might be taken as constant.

Some approaches in assessing small RNA-Seq signals take as a base line the total mapped reads in a library, and reads per million (RPM) of these are compared. Bedgraphs, used as input for COALISPR, are based on RPMs. Quantitative comparison between bedgraph signals from different experiments is, however, not always possible. When libraries differ significantly in their number of reads or in the case of considerable differences in the amount of background reads, numerical comparison does not seem to be reliable or trustworthy. Size selection of siRNAs cannot evade contamination by equally-sized break-down products of larger molecules like rRNA or tRNA which, due to their abundance, can be responsible for large (but not necessarily equally large) fractions of total read counts. Different RNA isolation methods, like size selection of small RNAs from total RNA vs. RNA-IP with Argonaute proteins, can lead to incomparable differences in co-purified background. This makes it difficult to quantify siRNA read-counts by normalisation to mapped library sizes, especially when yields of library preparations vary.

Experimental design decisions (inclusion of a spike-in RNA, or combining samples during cDNA synthesis) will contribute to obtain data that will be better comparable but will not address the above sketched difficulty in quantitative comparison posed by the low number of siRNA loci. Some bioinformaticians argue that the compositional (or relative) nature of count data requires that counts need to be assessed in relation to the mixture of reads in a sample. This aspect, the proportionality of counts, is inherent to the methods by which RNA-Seq libraries are generated (37, 38). Looking proportionally at RNA sequencing data by compositional data analysis (CoDa) is not generally done; most methods take the read counts as is and try to normalise these to some standard. To turn this into a valid approach for siRNAs is, however, not straightforward.

### Sourcing ncRNAs

Many abundant RNA molecules, like rRNAs and small ncRNAs, form common background signals in RNA-Seq experiments. Given that most of such molecules are not or only poorly annotated, one can treat the negative control data as a resource to get an impression of the population of ncRNAs in these cells. In the case of the *Cryptococcus* samples, we have been able to identify the full extent of the genes for 35S pre-rRNA, a series of tRNAs, some of which had not been annotated, SRP RNA (39) (Figure 7A) and – previously unidentified – U1 snRNA (Figure 7A, C), and other biologically important molecules like RNase P, or H/ACA and box C/D snoRNAs, for example U3, U14 or snR190, all important for processing of pre-rRNA (Figure 7A, D).

Like in other eukaryotes, cryptococcal snoRNAs are formed by transcription of snoRNA genes or are processed from introns that have been removed by splicing from precursors of protein-coding messages (40). For example, U14 is encoded by the 2^nd^ intron, snR190 by the 1^rst^ intron in *Argininosuccinate synthase* genes of *Cryptococcus* (Figure 7A) which seems conserved in related *Tremellales* (jelly fungi). Both snoRNAs are linked to different genes in more distant *Tremellomycetes* but appear in the same 5’ to 3’ order as first found in *S. cerevisiae* (41), where it is encoded on a bicistronic transcript. In some fungal species each snoRNA is expressed from a different locus.

Genomic organisation has diverged in evolution for other box C/D snoRNAs, like for snR51, snR41 and snR70. In *Saccharomyces* these snoRNAs are expressed as part of a cluster that is processed (snR41-snR70-snR51), while in *Tremellomycetes* snR41 and snR70 derive from introns of a non-coding transcript that in the first intron expresses another snoRNA. The homologue for snR51 was found in a separate unit.

Thus, in *Cryptococcus*, another source for snoRNAs is formed by non-coding transcripts that are spliced and where the sole conserved section is the intron containing the snoRNA (Figure 7B). As a corollary, many ncRNAs for *C. neoformans* have been annotated with respect to their exons, while only the precursor would have been functional by providing a (conserved) snoRNA; the spliced exons do not contain conserved sequences that could point to some biological function. Of course, versatile as this organism is, also transcripts for snoRNAs are seen where one snoRNA is intronic and the other derives from an exon, e.g. in the case of CNAG_12256 with snR13 (intron) and snR45-U13/U3 (exon). Further, snoRNAs can be split by introns, even when encoded in an intron of a pre-mRNA, for example snR50-snR40l.

In yeast and higher eukaryotes, acetylation of rRNA nucleotides by Kre33 (NAT10) is guided by box C/D snoRNAs, snR4, or snR45 (U13) (11). In *Tremellomycetes*, if Kre33 modifies the same cytosines in 18S rRNA, the role of snR45 could be regulated or implemented by two different snoRNAs (snR45-U13_U3a and snR45-II), each with the potential to provide a separate guide region for attachment to the 18S rRNA region near the putative substrate (the counterpart of C1773 in 18S rRNA of *S. cerevisiae*) (Supplementary Figure 4). SnoRNA snR45-U13_U3a has a homologue, U3b, and both these snoRNAs fit the Rfam signature RF00012 for the small nucleolar RNA U3 despite some changes in secondary structure: in *Tremellomycetes* the 5’ leader sequence upstream of the C-box in these snoRNAs differs from the leader in U3 of baker’s yeast where a well-conserved helical region confers base-pair interactions with 18S rRNA sequences that form an essential pseudo-knot in the mature rRNA (42). The extent in which these nucleotide interactions can be provided by the U3 snoRNAs of *Tremellomycetes* appears to vary (Supplementary Figure 4), leaving the possibility that additional events might be supported by these snoRNAs.

For some snoRNAs, for which no counterparts in other organisms were identified, no obvious base-pairing partner could be identified. For example snRcnag12093 (Figure 7B) is well-conserved among *Tremellomycetes* but putative targets were not obvious. Methylation involves dynamic, short-lived base-pair interactions which might be hampered in the case of some loci where long stretches of complementarity (>13 bp) between the guide of the snoRNA and pre-mRNAs was found nearby splice junctions or a branch-point. Provided these RNAs are able to interact, this could imply a side-effect of the presence of this snoRNA on storage or processing of splicing intermediates in *Cryptococcus*. In all, over 120 ncRNAs not previously annotated for *Cryptococcus* were identified and validated by phylogenetic analyses. All details are available with the documentation of COALISPR. Experimental confirmation will be needed to get definitive proof for functional relevance of the less conserved ncRNAs.

## CONCLUSION

By incorporating negative controls in bioinformatics analysis of high-throughput datasets we could easily identify meaningful signals and obtain count data without relying on a reference with annotated regions. By doing so, it was possible to assess the variability between libraries, which exemplified difficulties inherent to common practices when applied for quantitative comparison of small RNA-seq data. We demonstrated that coverage and abundance of ribosomal RNA fragments (rRFs) and other abundant ncRNAs are comparable between negative and positive datasets to such an extent that no biological meaning can be assigned to their co-purification with Argonaute proteins. In general, because the amplification steps during high-throughput sequencing elevate the levels of non-specific RNAs and intensify the background signals, the occurrence of rRFs or other abundant RNAs in co-purification studies require careful evaluation with respect to their biological significance; such observations might actually be highly ‘unlikely’ (1) and should be explicitly checked against sequencing data obtained for appropriate negative controls – even when the RNA input appears to be ‘clean’ – as shown here.

Although not providing information relevant for the biological system under study, negative datasets can help to annotate highly abundant RNAs and assess their roles. The RNA-Seq results for *Cryptococcus* species, combined with phylogenetic comparison, guided identification of a large number of snoRNAs and other ncRNAs present in these and related fungi. Many of these RNA molecules are processed from intronic sequences spliced from protein-coding pre-mRNA transcripts as found in higher eukaryotes. Novel is that some snoRNAs are derived from non-coding transcripts that are spliced but only carry sequences with significant evolutionary conservation within their introns. In these cases the precursor is functional, not the annotated spliced transcript.

At the basis for these insights was the Python application COALISPR we developed to process small RNA-Seq data for various organisms. By means of negative control data, the program provides a systematic approach to separate reads that might be biological relevant from those that form unspecific background. The program makes explicit what mostly remains hidden in experimental practicalities, the biologists’ technique to remove noise from signals.

## ACKNOWLEDGEMENTS

The author thanks Elizabeth H. Bayne for the opportunity to set up the RNAi project on *Cryptococcus deneoformans* in her lab, thereby gaining practical and theoretical knowledge on the workings of Argonaute proteins and small interfering RNAs in fungi (and the pitfalls analysing these). The author is very grateful to Sander Granneman for constructive criticism on early drafts of this essay, the discussions about programming in PYTHON and PANDAS, and teaching high-throughput analysis techniques. The author is indebted to Markus Hafner and Aishe A. Sarshad for kindly providing their mouse miRNA sequencing data which enabled re-analysis of rRFs and helped development of COALISPR.

## ADDENDUM

In their pre-print Taheraly et. al (43) described comparable findings for siRNAs targeting exons and identified similar (intronic) sn(o)RNAs in *Cryptococcus neoformans*.

