## Supplementary material for "*Bio-*informatics: Integrate negative controls to get the good data": SupplementaryData_rvn_2.pdf

Supplementary Tables. Mouse count data annotated and sorted.
Outputs of ANNOTATE scripts of COALISPR for unspecific reads (sheet S1, red) and specific reads (sheet S2, blue). Note that rRFs form the majority of counted reads in the unspecific dataset (row 1), on a par with the specific counts for a miRNA cluster on chromosome 7.

| | | | | |
|---|---|---|---|---|
| Positive samples: | A2n_1 | A2n_2 | A2t_1 | A2t_2 |
| Negative controls: | n_1 | n_2 | t_1 | t_2 |
| Annotated strands | plus | minus | | |

Supplementary Figure 1.

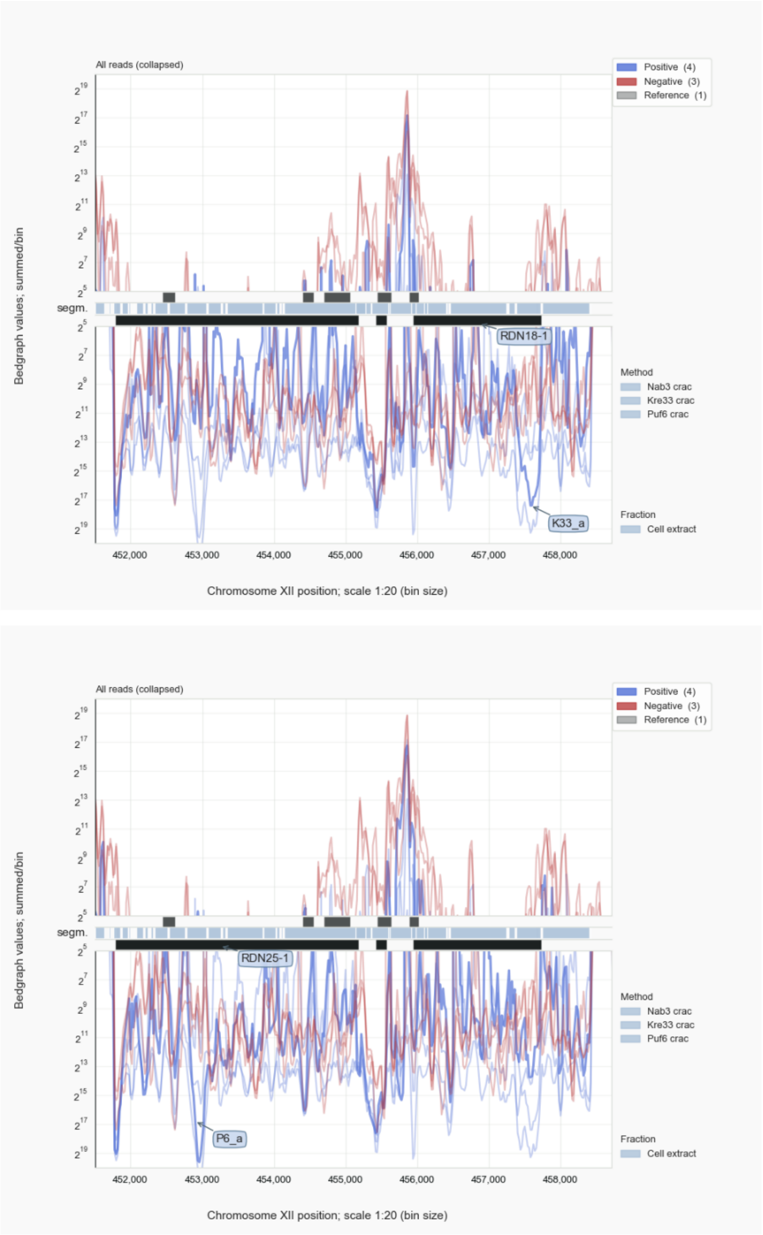


Supplementary Figure 1. UV crosslinking of RNA to pre-rRNA processing factors Kre33 and Puf6 in yeast *S. cerevisiae*. Kre33 crosslinks specifically to 18S rRNA regions (top); Puf6 binds helices in 25S rRNA (bottom)]. Nab3 (red) is the negative control. The relative increases of Puf6-signals for the Kre33-specific 18S segment (and vice versa for the Kre33 trace over the Puf6-region in 25S) is probably due to cross-contamination when these samples were prepared in parallel and amplified in the same PCR reactions. For other preparations, in which Kre33 and Puf6 crosslinks were separatedly assessed, these overlaps were not observed (data not shown). A kind of positive control for monitoring UV-crosslinked RNA fragments are single nt deletions (for CRAC) or mutations (for CLIP) that will be common for crosslinked uracil residues. These mutations reflect reverse trancription mistakes caused by remnants of amino acid adducts on a crosslinked residue.

Supplementary Figure 2.

In their seminal paper of 2013, Dumesic et al. (1) proposed that in *C. neoformans* H99 a pre-mRNA was either spliced or redirected to the RNAi machinery, which was based on the observation that many siRNAs mapped to introns. The annotation file they had used must have been a premature version, as almost all siRNAs actually align to exon sequences described in GTFs published since, including those for the original genomic sequence of this fungus described by Janbon et al. in 2014 (2).

For example, one of the signature genes in Dumesic et al. (1), CNAG_07721, has been re-annotated, with this chromosome 8 locus renamed to CNAG_03231 as was deduced from genome coordinates provided by Kim et al. (3) in their Supplementary Table S4 (sheet 3, entry 123). In *C. deneoformans* JEC21, siRNAs also align to exons (RvN, EHB data not shown) and for both fungi siRNAs crossing splice junctions were identified as shown in detail for *C. neoformans* in the section on intron-skipping siRNAs as part of the tutorials in the COALISPR documentation.

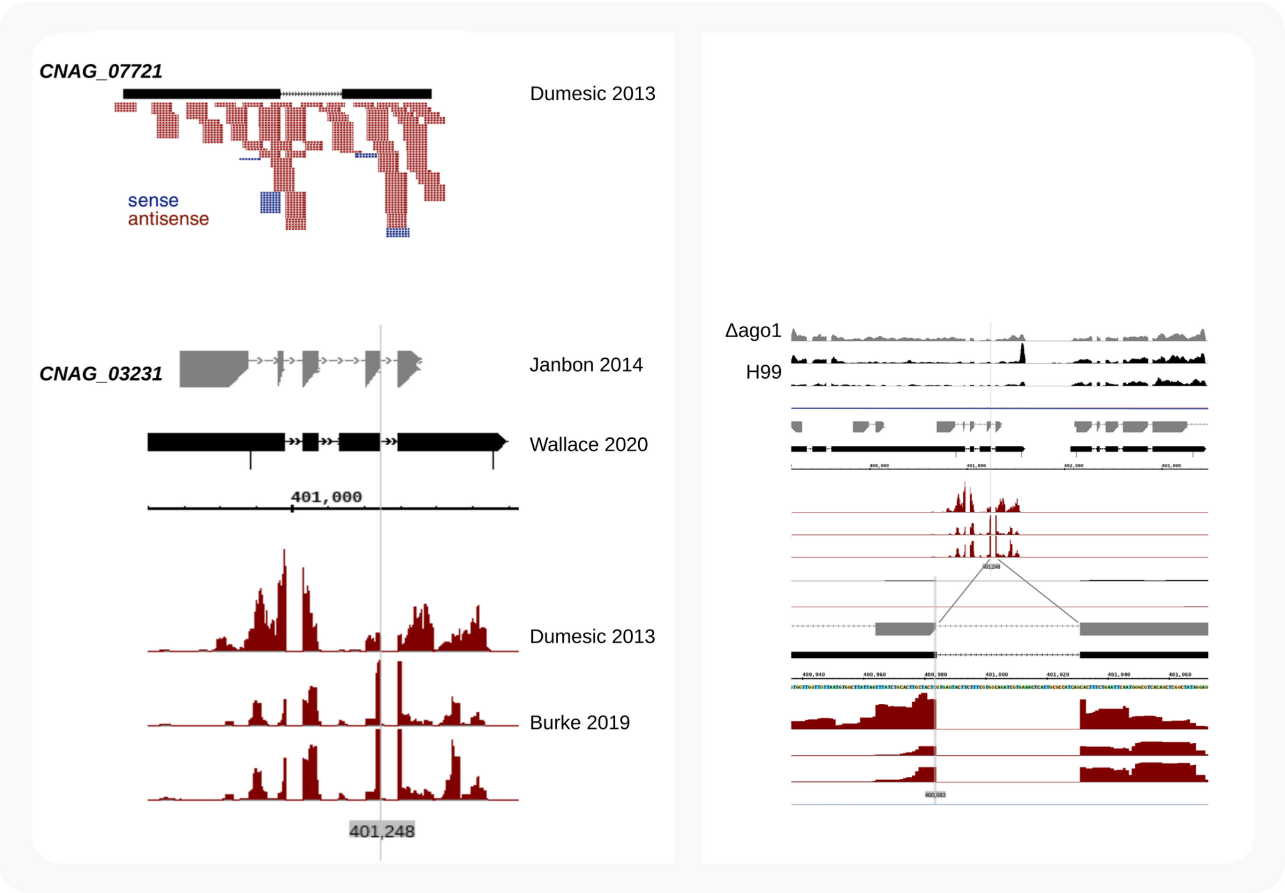


Supplementary Figure 2. *C. neoformans* siRNAs target exons, not introns. Left: Comparison of Fig. 1E from Dumesic et al. (1) and the locus for CNAG_03231, formerly named CNAG_07721, after mapping of wild type siRNAs from Dumesic et al. (1) and Burke et al. (4) by STAR as described in the Materials and Methods with annotations from Janbon et al. (2) and Wallace et al. (5). Right: Chromosme 8 context of CNAG_03231 and RNA-Seq by Wallace et al. (5) for two samples of wild type H99 (black) and for a strain with an Ago1 deletion (ΔAgo1; gray). Skipped introns have canonical splice site sequences (GU..AG; Bottom); siRNAs associated with Ago1 target the region of the transcript that encodes the C terminus of the protein.

Supplementary Figure 3.

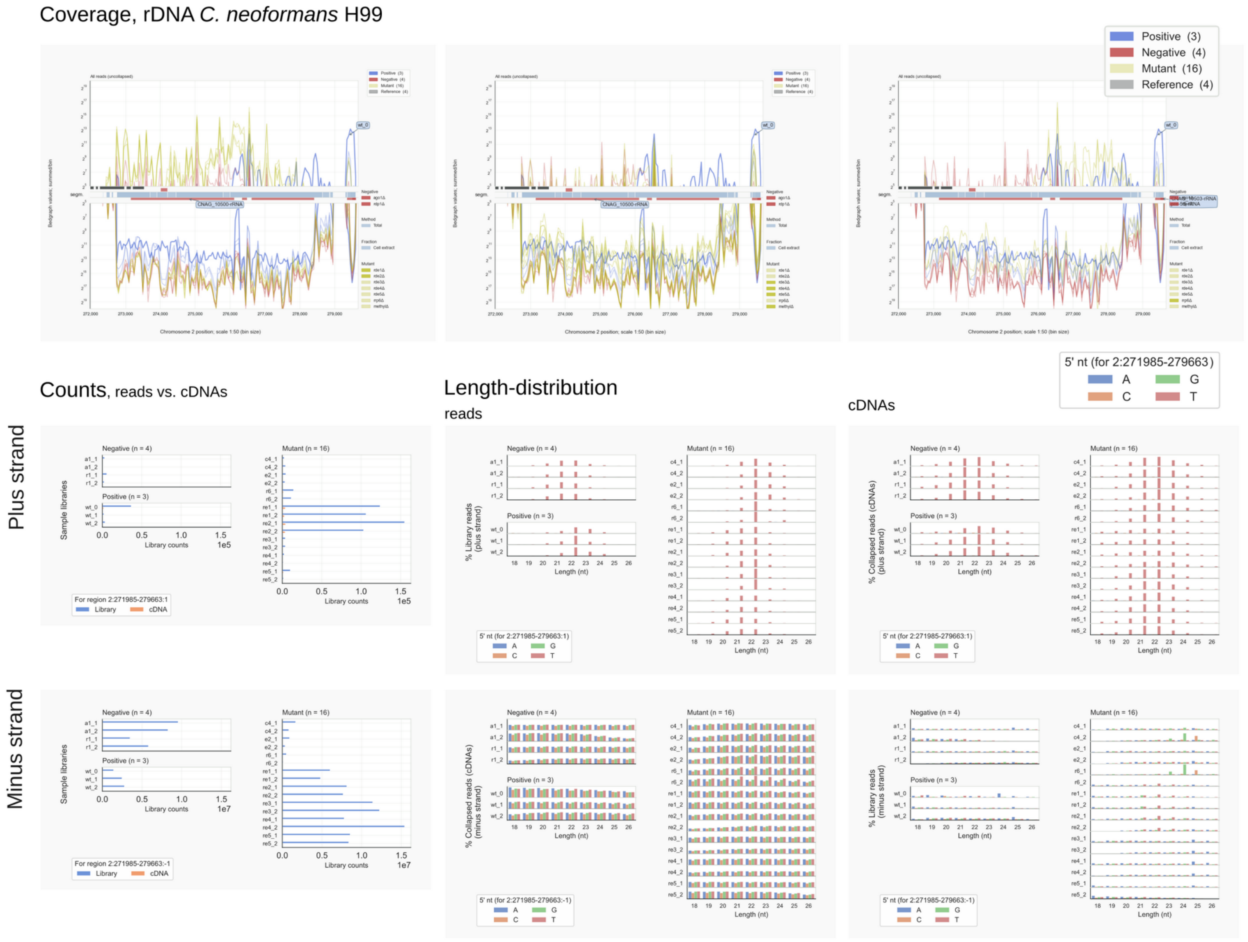


Supplementary Figure 3.. Analysis of *C. neoformans* RNAs  mapped to 2:271985-279663 for positive & negative controls, and rdeΔ, methylΔ or rrp6Δ mutants.

Reads representing rRFs are abundant in all samples of  Dumesic et al. (1) and Burke et al. (4) and cover the complete rDNA to a comparable extent in the controls (top right) and mutants. The pre-rRNA is transcribed from the bottom, minus strand. For rRFs no obvious difference is observed between all samples (bottom panel). In some clones, but not all, reads complementary to rRNA are formed in reasonable amounts that have the characteristic length and 5' nucleotide of siRNAs (middle panel). For the rde1Δ and rde2Δ strains (top left) these reads counter LSU (5.8S and 25S) rRNA regions, while in wild type (wt_0) or rrp6Δ (top right) the SSU (18S) rRNA region and flanking transcribed spacers are targeted. No effect is visible for strains without Rde3, Rde4 or Rde5 or in strains lacking DNA or histone methylation activity (top middle). Although a contamination with rDNA might have caused this, the signals are typical for the various strains and probably have another source of origin.

Supplementary Figure 4.

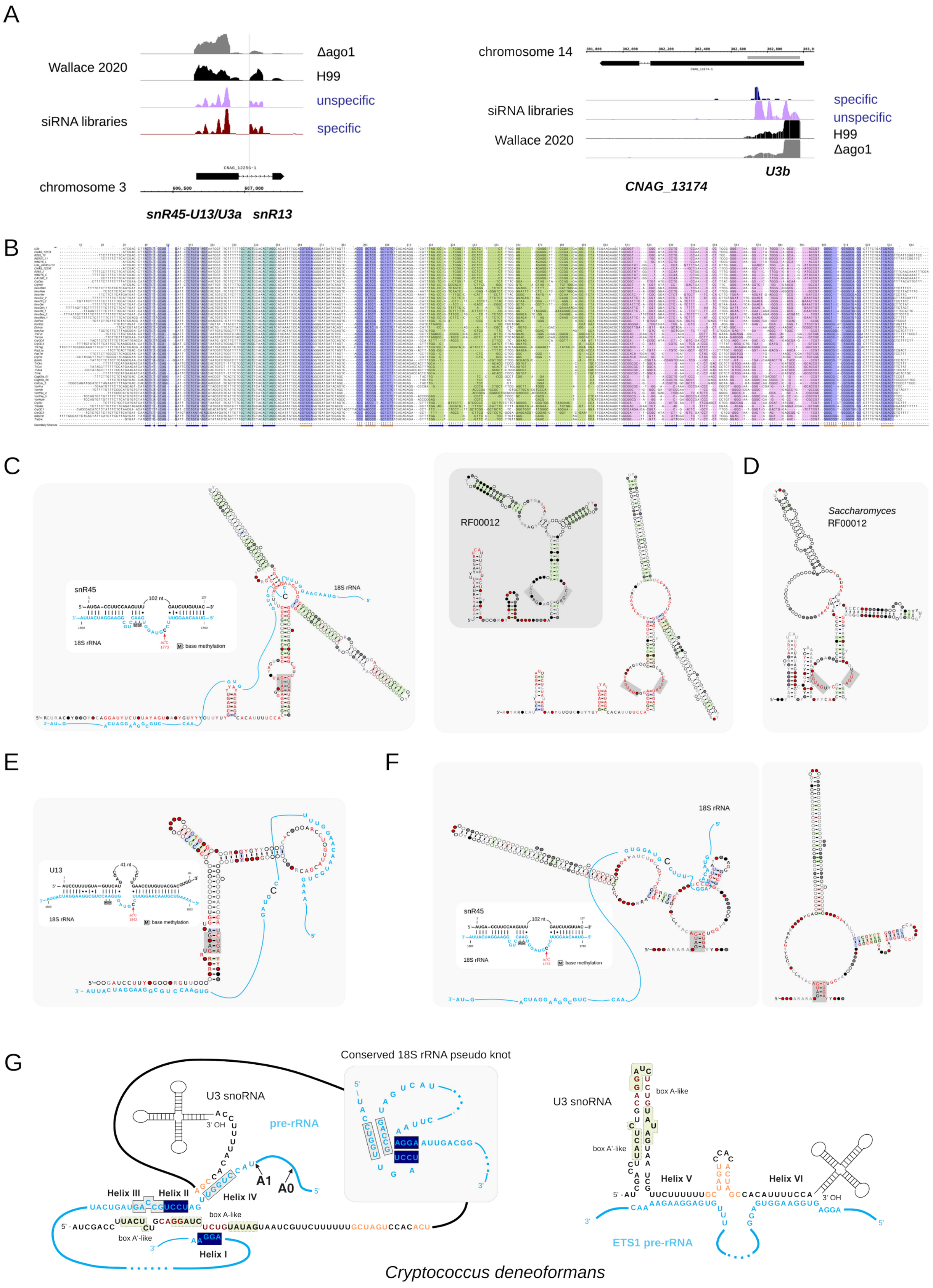

Supplementary Figure 4. U3 - and snR45-like snoRNAs in *Tremellomycetes*. In *Cryptococcus* two snoRNAs fitting the Rfam U3 signature (RF00012) are derived from exons (A) with homologues that adopt comparable secondary structures (B, C). The colours in the alignment (B) indicate stretches that are in the same helical region; the vertical blue line indicates the presence of an intron in some *Tremellomycetes*. Upstream and downstream sequences have been included for some sequences. R2R models depict U3 snoRNA folds for *Tremellomycetes* (C, right; with RF00012 in the inset) or *Saccharomyces* (RF01846) (D), and a putative snR45-like conformation (C, left). E: The mammalian counterpart of yeast snR45, U13 (6), can interact with 18S rRNA (blue) up - and downstream of the acetylated C (black) through two complementary regions, the 5' leader and a conserved guide in an internal loop. The *Tremellomycetes* U3 molecules, which deviate from yeast U3 by the reduced size of the conserved 5' stemloop with 18S-rRNA complementarity (C, right), might mimic the upstream interaction of snR45 (C, left). Another snoRNA in *Tremellomycetes*, snR45-II, could provide snR45-like interactions around and downstream of the methylated C (F, left). This snoRNA can adopt another conformation involving highly conserved nucleotides (F, right). Insets in C (left), E, and F (left) show base-pair interactions between 18S rRNA and yeast snR45 (C, F) and U13 (E) as proposed by Sharma et al. (6). Conserved Box C (RUGAUGA) and Box D (CUGA) motifs (gray background) form a helical segment in C (left), E, and F, but are unstructured in Rfam models of C (right) and D. G: Putative interactions between U3 snoRNA and pre-rRNA in *C. deneoformans* as based on Marmier-Gourrier et al. (7). The box A'- and box A-like elements differ from those in yeast; Helix I is less well supported (by three instead of four basepairs) because of changes in the nucleotide stretch linking the two conserved sections of box A.
